# Genome-wide CRISPR-Cas9 screen in *E. coli* identifies design rules for efficient targeting

**DOI:** 10.1101/308148

**Authors:** Belen Gutierrez, Jérôme Wong Ng, Lun Cui, Christophe Becavin, David Bikard

## Abstract

The main outcome of efficient CRISPR-Cas9 cleavage in the chromosome of bacteria is cell death. This can be conveniently used to eliminate specific genotypes from a mixed population of bacteria, which can be achieved both *in vitro*, e.g. to select mutants, or *in vivo* as an antimicrobial strategy. The efficiency with which Cas9 kills bacteria has been observed to be quite variable depending on the specific target sequence, but little is known about the sequence determinants and mechanisms involved. Here we performed a genome-wide screen of Cas9 cleavage in the chromosome of *E. coli* to determine the efficiency with which each guide RNA kills the cell. Surprisingly we observed a large-scale pattern where guides targeting some regions of the chromosome are more rapidly depleted than others. Unexpectedly, this pattern arises from the influence of degrading specific chromosomal regions on the copy number of the plasmid carrying the guide RNA library. After taking this effect into account, it is possible to train a neural network to predict Cas9 efficiency based on the target sequence. We show that our model learns different features than previous models trained on Eukaryotic CRISPR-Cas9 knockout libraries. Our results highlight the need for specific models to design efficient CRISPR-Cas9 tools in bacteria.

## Introduction

The Clustered Regularly Interspaced Short Palindromic Repeats (CRISPR)-associated protein 9 (Cas9) has been repurposed as a tool for a variety of applications including genome editing, transcriptional repression or activation, epigenetic modifications, chromosomal loci tagging and more (Hsu, Lander, and Zhang 2014). In most Eukaryotic cells, Cas9 breaks can be efficiently repaired by template-independent non-homologous end joining (NHEJ), which introduces small indels and can be conveniently used to knockout genes. Conversely, the main outcome of Cas9 cleavage in the chromosome of bacteria is cell death (Bikard et al. 2012, 2014; Citorik, Mimee, and Lu 2014; Gomaa et al. 2013). Most bacteria lack a NHEJ system, but even in species that do carry such repair pathway, it seems unable to efficiently repair Cas9 break under laboratory conditions (T. Xu et al. 2015; Bernheim et al. 2017). Bacteria mostly rely on homologous recombination to repair double strand breaks, which is made possible by the fact that several copies of the chromosome are present in the cell under most conditions (Dillingham and Kowalczykowski 2008; Ayora et al. 2011).

We recently showed that when Cas9 is guided to target a position in the chromosome, some guide RNAs appear to be less efficient than others. When guided by a weak guide, Cas9 will not cleave all copies of the chromosome simultaneously leaving a copy intact for repair and allowing cells to survive by entering a cycle of DNA cleavage and repair. However, a strong guide will lead to the simultaneous cleavage of all chromosome copies making repair through homologous recombination impossible and leading to cell death (Cui and Bikard 2016). This property can be used to select specific mutants or genotypes in mixed bacterial populations or even harnessed to engineer sequence-specific antimicrobials able to kill target antibiotic resistant or virulent bacteria (Bikard et al. 2014; Gomaa et al. 2013; Jiang et al. 2013; Citorik, Mimee, and Lu 2014). For all these applications it is critical to understand what makes some guide RNAs better at killing the cell than others. Cas9 cleavage is the result of a process which starts with the formation of the gRNA-Cas9 nucleoprotein complex. Target search then occurs through the recognition of a small protospacer adjacent motif (PAM), followed by R-loop formation through pairing of the guide RNA with the target strand (S. H. Sternberg et al. 2014; Anders et al. 2014; Jinek et al. 2012). Upon successful pairing a conformational shift in the Cas9 protein occurs bringing two catalytic domains, RuvC and HNH, in contact with the DNA and leading to the formation of a double strand break (Samuel H. Sternberg et al. 2015; Jinek et al. 2014). Each of these steps could be impacted by the sequence of the guide RNA, genomic context, DNA conformation and DNA modifications.

High-throughput screens performed in Eukaryotic systems have enabled to identify sequence features that determine knockout efficiency (Doench et al. 2014, 2016; H. Xu et al. 2015; Moreno-Mateos et al. 2015; Wang et al. 2014). However, the features that emerge in these models seem to be very different between experimental setups, and likely result in great part from constrains on the proper expression and stability of guide RNAs, rather than from their impact on Cas9 biochemical activity (Haeussler et al. 2016; Moreno-Mateos et al. 2015).

The goal of this study is to elucidate the genetic requirements for efficient CRISPR-Cas9 targeting in the bacterial chromosome. To this end, we performed a high-throughput screen in which we guide Cas9 to cleave ~92,000 different random positions around the chromosome of *E. coli* MG1655, followed by a “NGG” PAM. Upon Cas9 induction, guides that efficiently kill *E. coli* are expected to be rapidly depleted from the library, which can be measured through sequencing of the guide RNA library before and after induction of Cas9 expression. We observed large regions along the chromosome (~100 kb) in which the sgRNAs are rapidly depleted from the library while sgRNAs from other regions, are still present several hours after Cas9 induction. Surprisingly, this large-scale pattern is not correlated to cell death, as revealed by time-lapse microscopy. Indeed, cells immediately stop dividing, start to filament and then die after Cas9 induction regardless of the region targeted. We reveal that the pattern observed is rather due to differences in the copy number of the plasmid carrying the guide RNA library that arise after DNA cleavage in different regions of the chromosome. The loss of some genomic regions leads to an interruption of plasmid replication, while the plasmid can still replicate and increase its copy number after cleavage in other regions. The scale and shape of the pattern is determined by the extent of chromosomal DNA being degraded after Cas9 cleavage. After taking this interesting phenomenon into account, one can still observe differences in the efficiency of guide RNAs targeting the chromosome of *E. coli*. We built a neural network model able to predict this efficiency based on the guide sequence. Model previously build on Eukaryotic datasets have a limited predictive power on our data, highlighting the need to develop specific models of guide RNA activity in bacteria. Our model should directly aid in the design of guide RNA for genome editing and the development of CRISPR antimicrobials. We are making it available as guide RNA design tool at the following address: http://hub13.hosting.pasteur.fr:8080/CRISPRBact/.

## Results

### Large-scale depletion pattern of sgRNAs after Cas9 cleavage

We previously constructed a library containing 92,000 guides targeting random positions in the genome of *E. coli* followed by a “NGG” PAM motif (Cui et al. 2018). We introduced this library in *E. coli* MG1655 carrying Cas9 in the chromosome under the control of a Ptet promoter (strain LC-E19). Cas9 was induced and the library sequenced before induction and at different time points in order to determine the efficiency of killing of each guide (**Fig. 1a**). The experiment was performed in triplicates and the depletion of guides in the library was computed as the log2 transformed fold change of read counts normalized to a non-targeting guide RNA. When plotting the results along the chromosome of *E. coli*, we immediately noticed a large-scale pattern where guides targeting certain regions on a scale of ~100 Kbp are depleted from the population faster than guides targeting other regions. This pattern can be represented as the moving average of log2FC with a sliding window of 6kb, and is especially striking after 4H of induction (**Fig. 1b, c**).

**Figure 1.**
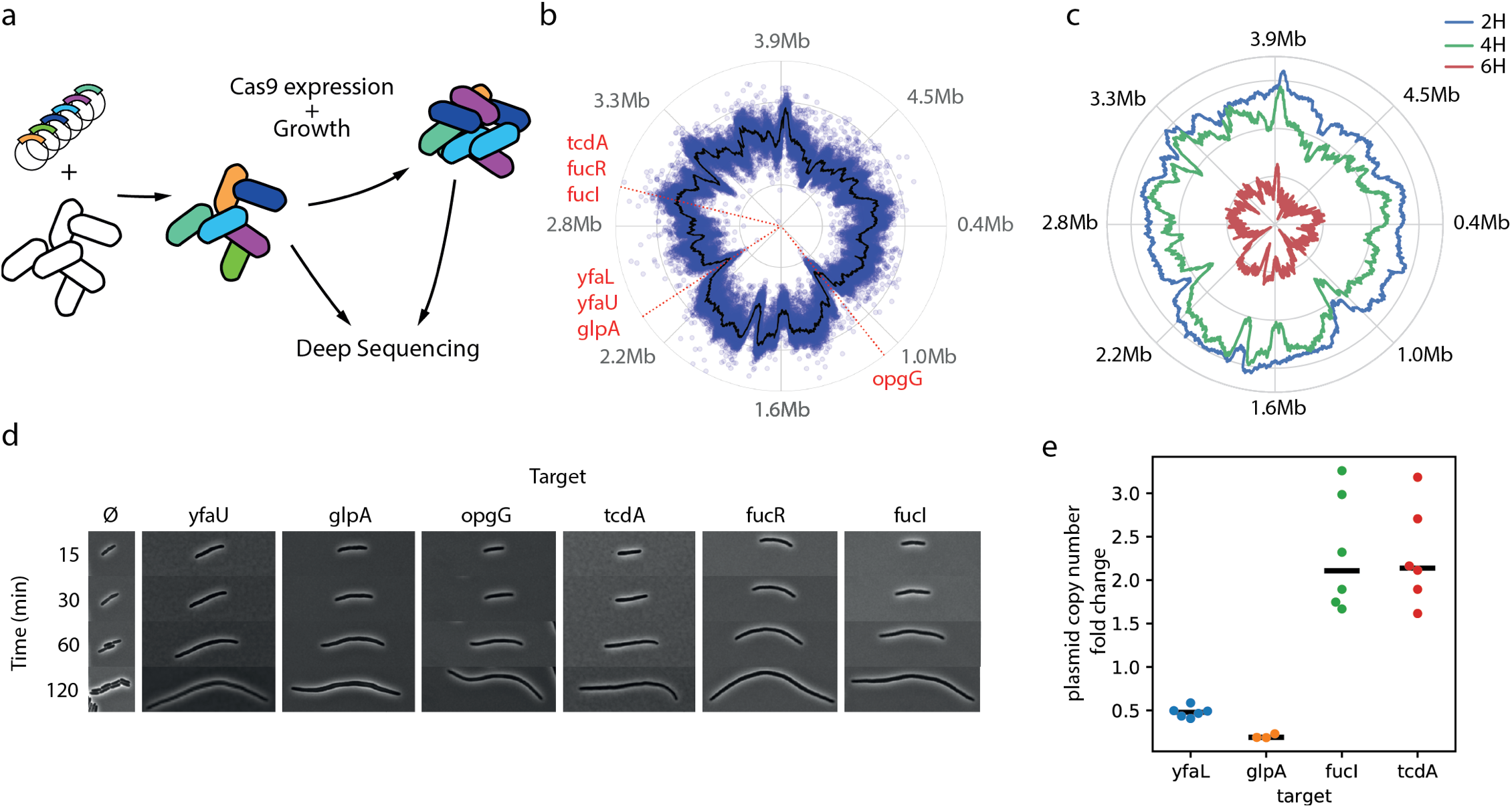
Genome-wide Cas9 killing screen reveals large-scale depletion pattern. a) A genome-wide library of guide RNA was introduced in *E. coli* strain LC-E19 carrying a *cas9* under the control of a Ptet promoter. Cells were grown in the presence of 1nM aTc and the guide RNA library sequenced before and after a few hours of induction. b) Scatter plot showing the log2FC of guides around the genome. The black line represents the moving average with a window size of 6kb (outer line of circle: log2F=2, centre of circle: log2F=-6). c) Moving average of guide RNA depletion around the genome after 2H, 4H and 6H of aTc induction. d) Time lapse microscopy after Cas9 induction in the presence of different guide RNAs. e) Fold change of plasmid copy number normalized to a non-targeting control as measured by qPCR. Points show independent biological replicates, the black bar shows the median.

We initially formulated the hypothesis that some chromosomal regions might be less accessible to Cas9 cleavage or more easily repaired than others. If this was the case, cells targeted in these regions might be able to survive longer and possibly keep dividing for a few generations after the induction of Cas9. To investigate whether the pattern observed through sequencing was indeed a measure of cell survival to Cas9 cleavage, we performed time-lapse microscopy experiments with guide RNA targeting different positions either in strongly depleted or weakly depleted regions. In all cases we observed that cells started to filament within minutes after Cas9 induction. Cells targeted by weakly depleted guides did not show a delayed or weaker response to Cas9 induction.

The number of reads obtained when sequencing the library can be seen as a measure of the abundance of plasmids carrying each guide RNA in the sample, which is typically used as a proxy of the number of cells carrying each guide in pooled CRISPR screen assays. Since the large-scale pattern observed did not seem to be linked to differences in cell survival, we reasoned that it might rather be due to differences in plasmid copy number (PCN). To test our hypothesis, we measured changes in PCN by qPCR and normalized to the number of copies of the chromosome as measured with qPCR probes distant from the cleavage position. We specifically investigated two regions: one centred on position 2.35 Mb in which guides are strongly depleted and one centred on position 2.95Mb in which guides are weakly depleted. We measured a decrease of PCN after Cas9 induction when targeting genes *yfaL* and *glpA* in the first region, and an increase of PCN when targeting genes *fucI* and *tcdA* in the second region (**Fig. 1e**). These results support the hypothesis that variations in the number of copies of the plasmid carrying the guide RNAs are responsible for the pattern observed.

### Degradation of chromosomal genes that control plasmid replication explains the large-scale pattern

We then sought to understand why targeting specific genomic regions affects plasmid copy number. We hypothesized that some regions might contain genes necessary for efficient plasmid replication. Targeting these regions with Cas9 will lead to the degradation of the target DNA and therefore genes in the target region will no longer be expressed. If our hypothesis is correct then it should be possible to restore plasmid replication by cloning genes in the target region on a plasmid that will not be targeted by Cas9. To test our hypothesis we cloned ~13.3 Kbp located in the central part of the peak located around 2.34Mbp in plasmid pBG7 (**Fig. 2a**). We then targeted different positions inside or just outside of the region and measured PCN after Cas9 induction. As expected, the copy number of the plasmid carrying the guide was higher in the presence of pBG7 than with a control empty vector (pBG10). When a different region in which guides are also strongly depleted was targeted (see target in *opgG*), pBG7 did not have an effect on the PCN of psgRNA (**Fig. 2b**). These results support our hypothesis that genes cloned on pBG7 are important for replication of the plasmid carrying the library and that the loss of these genes after Cas9 cleavage results in the interruption of plasmid replication. In order to identify the specific gene responsible for the variations in plasmid copy number, we independently cloned genes or operons present in the 13.3 Kbp region cloned on pBG7 (**Fig. 2a**). We only observed a substantial increase in PCN when the *nrdA-nrdB-yfaE* operon was cloned on the plasmid (**Fig. 2c**, see pBG15). Genes *nrdAB* encode a ribonucleoside diphosphate reductase involved in pyrimidine deoxyribonucleotides synthesis (Kolberg et al. 2004). The loss of these genes might lead to a decreased availability of deoxypyrimidines and consequently an inhibition of DNA replication. This hypothesis is also consistent with the fact that the level of *nrdA* mRNA strongly dropped 4H after Cas9 cleavage, consistently with the emergence of the pattern (**Supplementary Fig. 1**).

**Figure 2.**
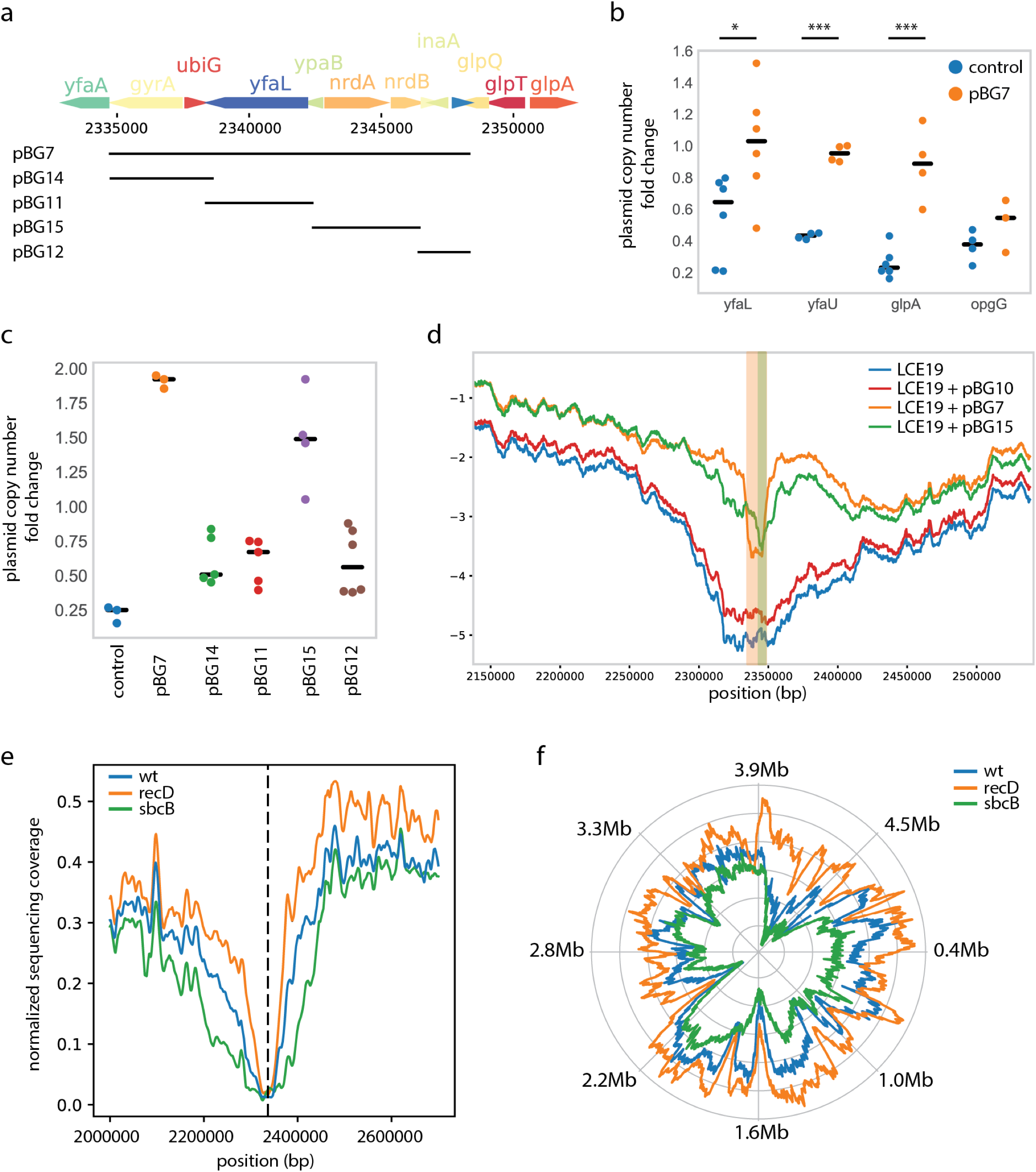
The large-scale pattern is explained by the degradation of chromosomal regions carrying genes necessary for replication. a) Genes at the centre of a depletion peak were cloned on a set of vectors. b) Fold change in the copy number of the plasmid carrying the guide RNA in the presence of plasmid pBG7 or the control empty vector pBG10, after cleavage with guides targeting genes *yfaL, yfaU, glpA* in the region, and *opgG* outside of the region. c) Fold change in the copy number of the plasmid carrying the guide RNA in the presence of various complementation plasmids and the control (pBG10), after 4H of induction with a guide targeting *glpA*. d) Moving average of guide RNA depletion in the genome-wide Cas9 killing screen performed in the presence of various complementation plasmids (window size of 6kb). e) Sequence coverage after 30min of Cas9 induction to cut position *yfaL* in strain LC-E19 (wt), a *ΔrecD* mutant and a *ΔsbcB* mutant (moving average with a window size of 10kb. f) Moving average of guide RNA depletion in the genome-wide Cas9 killing screen performed in *ΔrecD* and *ΔsbcB* mutants (window size of 6kb).

Finally, we performed the Cas9 genome-wide screen again in the presence of plasmid pBG7, pBG15 or the control empty vector. We observed a much weaker depletion of guides targeting the region located around 2.34Mbp in the presence of pBG7 and pBG15 than with the control vector (**Fig. 2d**). Interestingly guides in the library that targeted the region cloned on the complementation plasmid showed a stronger depletion than guides targeting just outside of this region (see Supplementary Fig. 2 for a zoom in the region). This is consistent with the fact that these guides will destroy both the chromosomal and plasmidic *nrdAB* operons, preventing rescue. As an internal control, a mutation was introduced in the PAM motif of one of the target positions carried by plasmid pBG7. As expected the guide targeting this position was much less depleted than surrounding guides (**Supplementary Fig. 2**).

### The shape of the large-scale pattern is determined by DNA degradation speed

Our results therefore indicate that the *nrdAB* genes might be responsible for the depletion of guides targeting a region of ~100 Kbp with guides further away from the genes being on average less depleted than guides closer to the genes. These results suggest that the extent of DNA degradation determines the shape of the depletion pattern. To investigate this in more details we programmed plasmid psgRNA to target gene *yfaL* (next to *nrdAB*) and sequenced the DNA extracted from strain LC-E19 after 30min of induction. Sequence coverage can be used as a measure of the abundance of DNA around the target region in the sample. DNA was degraded over ~50 Kbp away from the target in about half of the cells (**Fig. 2e**). The scale of DNA degradation matches that of the large-scale guide RNA depletion pattern consistently with our hypothesis.

If DNA degradation determines the shape of the guide RNA depletion pattern, mutants of genes involved in DNA degradation might change it. We repeated the whole genome Cas9 cleavage screen in *ΔrecD* and *ΔsbcB* mutants. The pattern was indeed strongly affected in these mutants with broader peaks and a stronger overall depletion of guides in the *ΔsbcB* mutant and narrower peaks with a weaker overall depletion of guides in the *ΔrecD* mutant (**Fig. 2f**). Consistently with this observation, DNA degradation was faster in the *ΔsbcB* mutant and slower in the *ΔrecD* mutant. Altogether, our results demonstrate that the large-scale pattern of guide RNA depletion originates from the effect of degrading some chromosomal regions on plasmid replication and copy number.

### A neural network model enables to predict guide RNA efficiency

The large-scale pattern described above accounts for an important part of the variation in the effect of guides, but we also observed a reproducible variability between guides targeting nearby positions. In order to eliminate the large-scale effect of plasmid copy number, we computed for each guide in the library an activity score as the difference between log2FC and the mean log2FC of all the guides within 6 kbp. The sgRNA activities were extremely consistent across experiments, showing on average Pearson correlation between replicates of 0.92 (Supplementary Fig 3). We then sought to model guide activity based on the one-hot-encoded target sequence (4 bases upstream, target, PAM and 3 bases downstream). A simple linear model with L1 regularization was able to predict guide activity in cross-validation with a Pearson correlation of 0.58. The coefficient of the features in the regression are plotted as a heat map in Figure 3a. The model uses positions in the guide RNA but not the surrounding region in the target. Interestingly bases at the 5’end of the guide RNA impacted the model predictions the most. The presence of thymidine and to lesser extent adenines seems to reduce the efficiency of the guides, while guanines and to a lesser extent cytidines increase it. This observation differs from previous observations in mammalian cells where the PAM-proximal region was more important than the distal region (H. Xu et al. 2015; Moreno-Mateos et al. 2015; Doench et al. 2014). Conversely, the presence of a guanine at the last position of the guide has a positive effect in our model, which was also consistently reported in previous studies.

**Figure 3.**
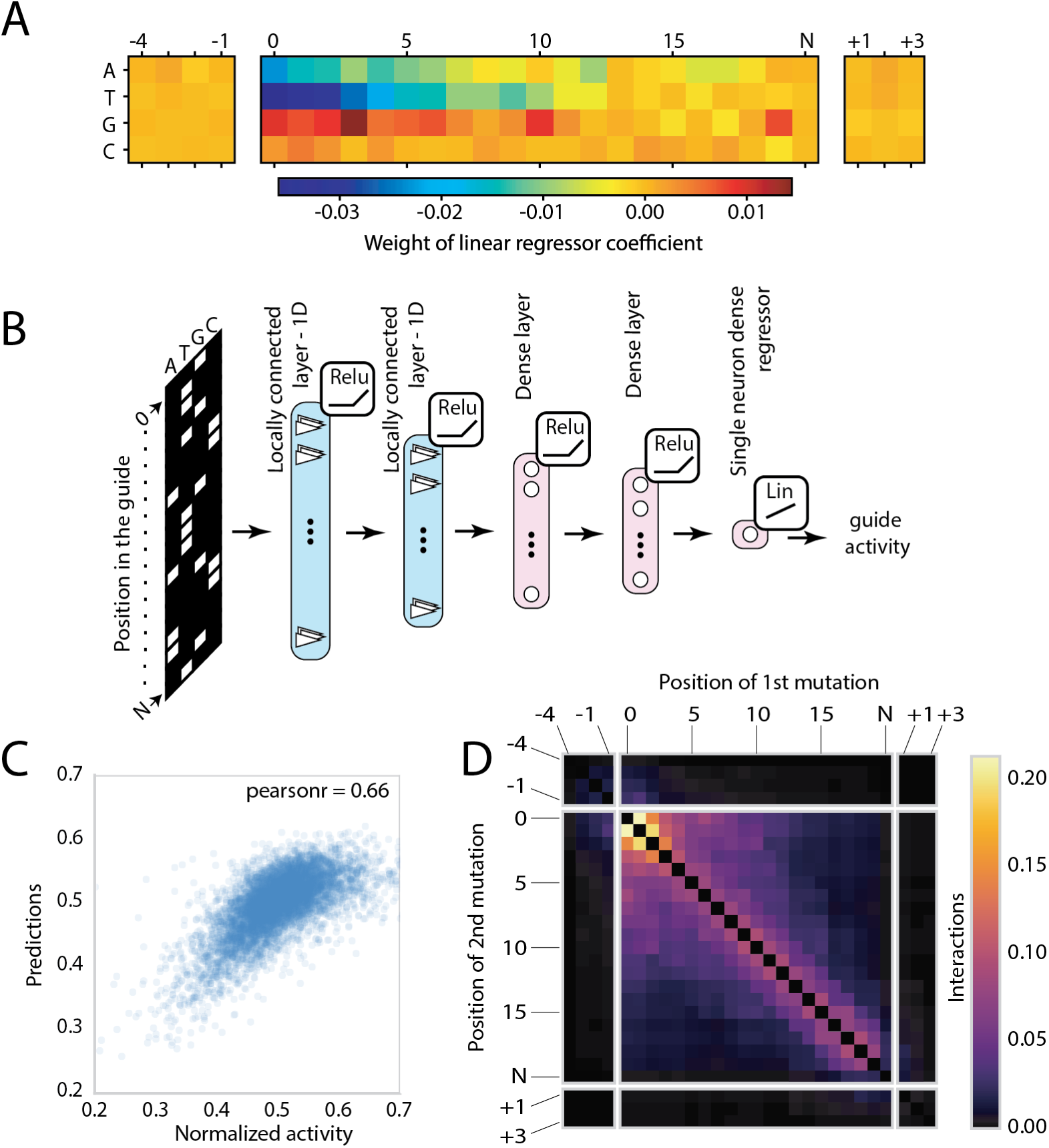
Predicting guide RNA efficiency. (A) A linear model with L1 regularization was trained to predict guide activity using the primary sequence as the only input feature. The heat map shows the coefficients attributed to each base. Position 0 is the first base of the guide, position N refers to the undetermined base of the PAM (NGG), position +1 refers to the first base after the PAM. (B) Architecture of the locally connected neural network trained to predict guide activity from the one-hot-encoded sequence. (C) Predictions of the model plotted as a function of the measured guide activity on a held-out test set. (D) Level of interaction between positions along the target that is seen by the model.

The ability of a simple linear model to obtain good predictions suggests that the determinants of guide efficiency in *E. coli* are quite simple and can for the most part be inferred from the specific bases present at each position. This model is however not able to encode more complex interactions between positions. The large amount of data collected in this experiments enables to train neural network models which have recently been successfully employed to model sequence data (Kim et al. 2018; Alipanahi et al. 2015). The dataset was split into a training, validation and test set. We first used a locally connected neural network to predict guide activity using an arbitrary 60nt window around the target (Figure 3b). The resulting model achieved a Pearson correlation coefficient of 0.66 on the held-out test set (Spearman-r=0.64). In order to investigate the sequence features used by the model to make its predictions we generated a set of 1000 random sequences and measured the impact on the model predictions of mutating each position to all possible bases. The results are consistent with the linear regression model, showing the importance of the 5’end of the guide, while giving little to no importance to positions outside of the 20nt target (Supplementary Figure 4). We thus restricted the inputs of the model to target sequence, 4 bases upstream and 3 bases downstream of the PAM. This simpler model still achieved a Pearson correlation of 0.66 on the test set while reducing overfitting on the training set (Figure 3c).

In order to investigate the interactions identified by the model we further mutated each pair of position to all possible bases for a set of 200 random sequences. We then compared the effect of a double mutation and the sum of the effect the two single mutations. It appears clearly that bases in the guide interact with their direct neighbours, and to a lesser extent with their second and even third neighbours (Figure 3d). Interestingly positions in the first half of the guide seem to interact with each other on a long range, an effect not observed in the PAM-proximal region.

### Comparison to other models

We then compared our model to two of the most widely used models for on-target activity prediction. Doench and colleagues modeled the activity of guide RNAs expressed from a U6 promoter on a DNA vector in human cells (Doench et al. 2016), while Moreno-Mateos and colleagues modeled the activity of guide RNAs expressed *in vitro* from a T7 promoter and injected into zebrafish embryo (Moreno-Mateos et al. 2015).

**Figure 4.**
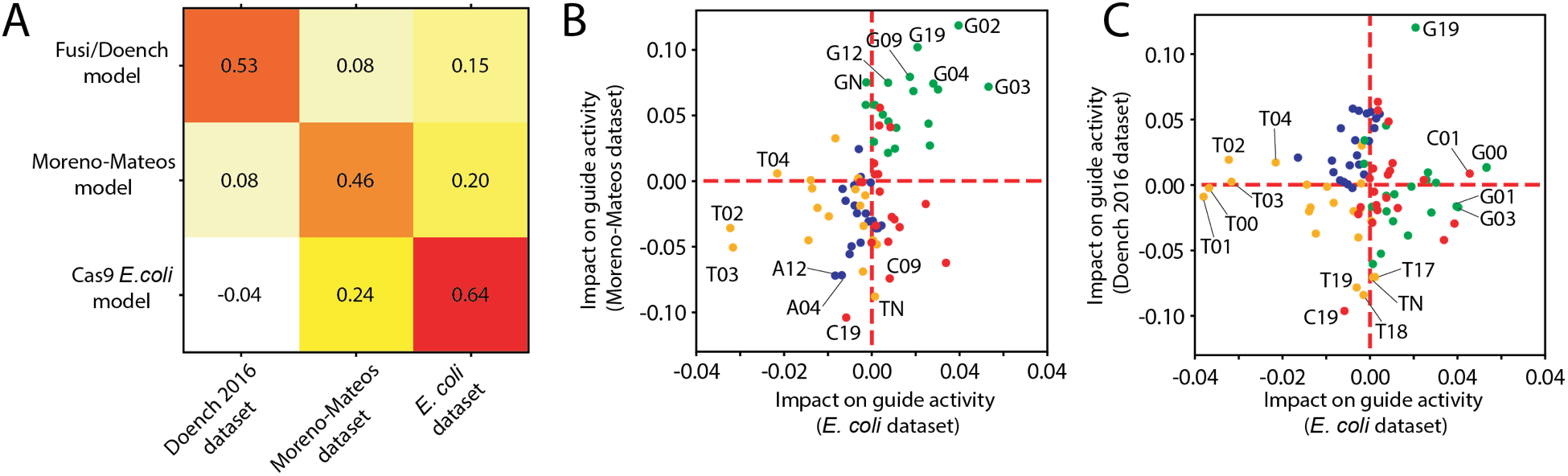
Comparision between models. (A) Heat map of Spearman correlation coefficient between activity scores predicted by models and datasets. The score given for the performance of models on their own training set corresponds to the performances on a test set in cross-validation. The other correlations are computed on the whole dataset. (B, C) Comparison of the impact of specific bases along the guide sequence between the different datasets, i.e. the difference between the average activity of guides that have this feature and the average activity of guides that do not have it. The following color code is used for bases (Adenine: blue, Thymine: yellow, Guanine: green, Cytosine: red).

A previous report showed that these two models make very different predictions suggesting that the determinants of guide efficiency strongly depend on the exact experimental setup (Haeussler et al. 2016). When applied to our data, the model of Moreno-Mateos provides some predictive power, and conversely our model shows some predictive power on the data from Moreno-Mateos. The model of Doench performs weak predictions on the *E. coli* and zebrafish data. In order to shed light on the specific sequence features that matter in these different datasets, we plotted the impact of having specific bases at each position along the guide RNA, i.e. the difference between the average activity of guides that have this feature and the average activity of guides that do not have it. This analysis reveals that many of the features that impact the activity of guides in *E. coli* also impact the activity of guides in the zebrafish dataset. Note that because of experimental constrains, the dataset of Moreno-Mateos only includes guides whose first two bases are guanines. In our dataset these two bases are the most important features determining guide activity, but their effect is unknown in the experimental setup of Moreno-Mateos.

## Discussion

Over the last few years numerous studies have led to a detailed understanding of Cas9 target search mechanism, structure, and biochemical activity (Gasiunas et al. 2012; Jinek et al. 2012; S. H. Sternberg et al. 2014; Jinek et al. 2014; Szczelkun et al. 2014; Anders et al. 2014; Samuel H. Sternberg et al. 2015). Despite the impressive amount of knowledge gathered, little is known about how the guide RNA sequence affects the efficiency of Cas9 targeting. Several groups have performed high-throughput screens in eukaryotic cells to measure the activity of thousands of guide RNAs, enabling the development of algorithms that can predict the efficiency with which a guide RNA can knockout a gene (Doench et al. 2014; Moreno-Mateos et al. 2015; H. Xu et al. 2015; Doench et al. 2016). Strikingly, these algorithms tend to perform well on some datasets and very poorly on others (Haeussler et al. 2016; Labuhn et al. 2018). Some of these differences have been attributed to the specificities of the experimental system and whether the guide RNA is transcribed *in vitro* from a T7 promoter followed by injection in cells, or *in vivo* from a U6 promoter on a DNA vector (Haeussler et al. 2016). For instance, assays based on DNA vectors consistently observed a bias against uracil at the 3’end of the guide RNA sequence, which is due to the propensity of the RNA polymerase III to terminate transcription at uridine-rich sequences (Doench et al. 2014; Wang et al. 2014; Wu et al. 2014; Moreno-Mateos et al. 2015). These results argue for the necessity to develop guide RNA design tools that are specific to each experimental setup.

Here we measured guide RNA depletion after cleavage in the chromosome of *E. coli*, enabling the investigation of features that affect Cas9 targeting in a completely different experimental setup than previous reports, and with a lot more data points. In this assay efficient guides kill the bacteria and are rapidly depleted. The most striking feature that explained differences between guide RNAs was their position along the chromosome. Studies in Eukaryotic systems have shown the importance of chromatin structure and accessibility on Cas9 targeting efficiency (Kuscu et al. 2014; Wu et al. 2014; Horlbeck et al. 2016; Chen et al. 2016). However, the pattern of guide RNA depletion that we observed here did not match the known structure of the *E. coli* chromosome (Lioy et al. 2018). Rather, we were able to demonstrate that this large-scale pattern emerges from changes in plasmid copy number that occur after the degradation of specific chromosome regions after Cas9 cleavage. The description of this artefact should serve as a cautionary tale for any pooled genetic screen in which the readout is made by sequencing a guide RNA or any barcode on a plasmid. Note that Cas9 might still be able to kill *E. coli* more or less efficiently depending on which region of the chromosome is targeted, but a different experimental design will have to be used to investigate this question.

After taking the large-scale depletion pattern into account we could investigate the effect of the guide RNA sequence on the ability of Cas9 to kill *E. coli*. Our results highlighted the positive effect of a high GC content at the 5’ end of the guide RNA, and in particular the negative effect of thymidines in this region. Binding of the 5’ end to the target sequence is required for a conformational shift to occur in Cas9 which brings the RuvC and HNH catalytic domains in contact with the target DNA (Samuel H. Sternberg et al. 2015; Jinek et al. 2014). This binding might be disfavored by a high thymidine content, making cleavage less efficient. Another possible explanation is that guide RNA transcription might be imperfect for some sequences. It was previously reported that the sequence around the transcriptional start site (TSS) can impact the frequency at which the polymerase will initiate transcription at different positions (Vvedenskaya et al. 2015). In particular, thymidine-rich sequences where shown to favor initiation further away from the −10 element, which in our case would lead to the formation of truncated guide RNAs. The same study also highlighted the fact that poly-T and poly-A stretches after the TSS promote slippage synthesis, which could conversely lead to the formation of longer guide RNAs. Finally, the sequence at the 5’ end of the guide might impact its stability *in vivo*. Moreno-Mateos measured an increased stability and activity of guides with a higher guanine content, and provided evidence that these guides might form G-quadruplexes protecting them against 5’-directed exonucleases. This mechanism could also be at play in our experiments.

Altogether, our results provide insight into the features that determine the ability of Cas9 to efficiently kill *E. coli*. We provide a model able to predict the most efficient guides with a high accuracy that should be useful for the design of effective guides in genome editing experiments as well as for applications of CRISPR as antimicrobials (Jiang et al. 2013; Cui and Bikard 2016; Bikard et al. 2014; Citorik, Mimee, and Lu 2014). This model is made available online: http://hub13.hosting.pasteur.fr:8080/CRISPRBact/

## Methods

### Bacterial strains and media

*E. coli* strains were grown in Luria-Bertani (LB) broth or LB Agar 1.5% as solid medium. Whenever applicable, media was supplemented with chloramphenicol (20 μg ml^−1^), carbenicillin (100 μg ml^−1^) or kanamycin (50 μg ml^−1^) to select or ensure the maintenance of the plasmids. Lower concentration of kanamycin (20 μg ml^−1^) was used to select for the integration of vectors in the chromosome. All the strains modifications derived from *E. coli* MG1655. *E. coli* strain DH5α or MG1655 were used as transformation recipients.

### Plasmids and *E. coli* strains construction

Strain LC-E19 was constructed using the pOSIP system (St-Pierre et al. 2013) and the backbones were removed using the pE-FLP plasmid(St-Pierre et al. 2013). The Ptet-Cas9 expression cassette was integrated at the HK022 *attB* and a *gfp* (Green Fluorescent Protein) reporter gene under the control the *sulA* promoter was integrated at λ *attB* to monitor SOS response. Genes *recD* and *sbcB* were deleted from strain LC-E19 using the lambda red recombineering strategy (Sharan et al. 2009). Plasmid pKD4 was used as a template to generate linear DNA fragments via polymerase chain reaction (PCR) followed by electroporation into strain LC-E19 carrying plasmid pKOBEG-A (Chaveroche, Ghigo, and d’ Enfert 2000). Colonies resistant to kanamycin were selected and the resistance gene was then removed using plasmid pE-FLP. The constructions were verified by PCR and sequencing. All the bacterial strains used in this study are listed in Supplementary Table 1 and the primers used for strain construction in Supplementary Table 2.

Fragments for plasmid constructions were generated by PCR or restriction digestion and assembled through Gibson assembly (Gibson et al. 2009). Novel guide RNAs were cloned into plasmid psgRNA by golden gate assembly (Engler, Kandzia, and Marillonnet 2008). A list of plasmids is provided as Supplementary Table 3, and primers used in plasmid construction as Supplementary Table 4. A list of guide RNA used in the study is provided as Supplementary Table 5.

### Genome-wide Cas9 screen

The data shown in figure 1 was generated as follow. The guide RNA library carried on plasmid psgRNA was electroporated into LC-E19, plated in LB with kanamycin 50 μg/ml and incubated at 37°C for 4h. An estimated number of 10^7^ clones were recovered, pooled in 10ml LB and stored as 1ml aliquots in DMSO 10% at −80°C. To perform the assay, 1ml of frozen cells were thawed in 400ml of LB with kanamycin 50 μg/ml and cultivated at 37°C, 190 RMP until they reached early-exponential phase (OD600 ≈0.25). Then, aTc 1nM was added and cells were recovered at different time points (0h, 2h, 4h and 6h). Plasmids were extracted from 50 ml of culture using the NucleoSpin Plasmid kit (Macherey-Nagel, Duren, Germany). The whole assay was performed in triplicate.

During the course of the study we realized that leaky expression of Cas9 in strain LCE-19 might lead to the introduction of biases in the library as clones that mutate the CRISPR-Cas9 system or the target might be selected before induction with aTc. To avoid this problem, we used modified experimental design for the data shown in figure 2 and figure 3. Plasmid psgRNA carries a cos site enabling its packaging in phage lambda capsids. The library was transformed in strain CY2120 which carries a temperature sensitive lysogenic lambda prophage with its cos site deleted (Cronan 2013). Upon induction at 42°C, lambda capsids are produced and the psgRNA packaged. Cosmid particles can then be purified as described previously (Cronan 2013). Briefly, strain CY2120 carrying psgRNA was diluted 100-fold from an overnight culture in LB supplemented with 50 mM Tris-HCl buffer (pH 7.5) and grown at 30°C until OD600 ≈ 0.5. The culture was then incubated at 42°C for 20 minutes and then at 37°C for 4 hours. Cells were harvested by centrifugation at 4000g for 5 minutes, washed in lambda dilution buffer (Tris-HCL pH 7.5 20mM, NaCl 0.1M, MgSO4 10mM) with 1/20 of the culture volume, centrifuged and re-suspended again in the same volume of lambda dilution buffer. To purify the cosmid particles chloroform was added (20 vol%), samples vortexed for 15 seconds and incubated at 37°C for 15 minutes with shaking. Cosmids were harvested after centrifugation at 12,000g for 1.5 minutes.

This cosmid library was then used to transduce strain LC-E19 induced to produce Cas9 as follow. An overnight culture of strain LC-E19 was diluted 100-fold in fresh LB, 0,2% arabinose and aTc 1nM and grown until OD_600_ ≈ 0.6. The psgRNA library was transduced at a MOI (Multiplicity of Infection) of 0.2 and incubated at room temperature for 30 minutes. Cells were then grown 4h at 37°C, 190 RPM with kanamycin 50 μg/ml to inhibit the growth of cells that were not transduced. Cells were recovered and plasmids extracted from 15 ml of culture using NucleoSpin Plasmid kit (Macherey-Nagel, Duren, Germany). Transduction in strain MG1655, which does not carry Cas9, was used as a control.

To measure the relative abundance of guide RNAs in each sample, the library of guide RNA was sequenced following the method described by Lun Cui *et al*. (Cui et al. 2018). Two nested PCR reactions were used to generate the sequencing library with primers described in Supplementary Table 6. The 1^st^ PCR adds the 1^st^ index. The 2^nd^ PCR adds the 2^nd^ index and flow cells attachment sequences. Sequencing is then performed using primer LC609 as a custom read 1 primer. Custom index primers were also used: LC499 reads index 1 and LC610 reads index 2. Sequencing was performed on a NextSeq 500 benchtop sequencer. The first 2 sequencing cycles read bases common to all clusters and were set as dark cycles, followed by 20 cycles corresponding to the guide RNA.

We report here the log2 transformed fold change in the number of reads obtained for each guide RNA, normalized by the total number of reads obtained for the sample and the number of reads obtained for a control guide RNA which does not have a target position in the genome of *E. coli* MG1655. Reads were counted only if they showed a perfect match to the sequence of a guide in the library.

### Determination of plasmid copy numbers

Pre-cultures of LCE-19 carrying a plasmid with a single guide RNA were incubated at 37°C in triplicate at 190 RPM shaking overnight. The appropriate antibiotics (kanamycin 50 μg/ml or chloramphenicol 20 μg/ml) were used to maintain the plasmids. Pre-cultures were diluted 100-fold in fresh LB and grown in the same conditions until they reached early-exponential phase (OD_600_ ≈ 0.25). Cas9 expression was then induced with aTc 1nM. Cells were recovered before and after 4 hours of induction. Total DNA was extracted using the Wizard Genomic DNA Purification Kit (Promega, Madison, USA). Quantitative Polymerase Chain Reaction (qPCR) was carried out in triplicate for each extraction using the FastStart Essential DNA Green Master (Roche Diagnostics, Mannheim, Germany) in accordance with the manufacturer’s specifications in a LightCycle 96 system (Roche Diagnostics). The plasmid copy number was determined as described by San Millan *et al*. (San Millan, Heilbron, and Maclean 2013). The efficiencies of the reactions were calculated from a standard curve generated by performing qPCR with five 10-fold dilutions of template DNA in triplicate (10 ng/μl-1pg/μl). Primers BG114 and BG115 were used to perform the qPCR with a concentration of template DNA of 0.1 ng/μl. To determine the average PCN per chromosome, we used primers BG116 and BG117 to amplified gene *rpoB*. Primers are listed in Supplementary table 7. The amplification conditions were: initial denaturation for 10 min at 95°C, followed by 45 cycles of 10 s at 95°C, 20 s at 55°C and 10 s at 72°C. The following formula described by San Millan *et al*. (San Millan, Heilbron, and Maclean 2013) was used:

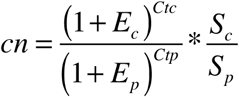

where *cn* is the plasmid copy number per chromosome, *S_c_* and *S_p_* are the sizes of the chromosomal and plasmid amplicons (in bp), *E_c_* and *E_p_* are the efficiencies of the chromosomal and plasmid qPCRs (relative to 1), and *Ctc* and *Ctp* are the threshold cycles of the chromosomal and plasmid reactions, respectively. Fold change values in plasmid copy number after 4H of Cas9 induction and normalized to a non-targeting control are reported.

### Quantification of gene expression

Overnight cultures were diluted 100-fold and grown until they reached early-exponential phase (OD_600_ ≈ 0.25). Then aTc (1nM) was added and cells incubated for 4h. RNA was extracted from 10 ml of culture at OD_600_ ≈ 0.25, and from 2ml after 4h of induction using Direct-zol™ RNA Miniprep kit (Zymo Research). All the RNA samples were first treated with DNase (Turbo DNase free kit, Ambion) and then 1 μg of RNA for each sample was reverse transcribed into cDNA using the Transcriptor First strand cDNA synthesis Kit (Roche). qPCR was performed using 1 μL of the reverse transcription reaction with the Faststart essential DNA green master mix (Roche) in a LightCycle 96 (Roche). Probes and PCR primers are listed in Supplementary Table 7. Relative gene expression was computed using the ΔΔCq method (Schmittgen and Livak 2008).

### Quantification of DNA degradation after Cas9 cleavage

Strain LC-E19 carrying plasmid psgRNA programmed to target gene *yfaL* was diluted 100x from on overnight culture and grown until OD_600_ ≈ 0.25. Cas9 expression was induced by addition of aTc 1nM followed by 30min of incubation at 37°C with shaking. Total DNA was extracted using the Wizard Genomic DNA Purification Kit (Promega, Madison, USA) and fragmented with a Covaris E220 ultrasonicator. Sequencing libraries were prepared using the NEXTflex PCR-free DNA-Seq kit (Bioo Scientific Corporation), and sequenced on a MiSeq v3 PE300 flowcell. Sequencing reads were mapped to the genome of E. coli LC-E19 using bowtie2 (Langmead and Salzberg 2012) and sequence coverage computed with samtools (Li et al. 2009). The plot of figure 2e shows a moving average of the coverage normalized between 0 and 1, with a window of 10kb.

### Microscopy

LC-E19 cells were transformed with plasmid psgRNA carrying different guides. Overnight cultures were diluted 100-fold in fresh LB and incubated at 37°C for 1h. Cas9 expression was induced with aTc 1nM and cells transferred to LB pads with 1% UltraPure™ agarose (Invitrogen), kanamycin 50 μg/ml and aTc 1nM. Cells were imaged using an inverted microscope (TI-E, Nikon Inc.) equipped with a 100x phase contrast objective (CFI PlanApo Lambda DM100x 1.4NA, Nikon Inc.). Images were taken every 5 min during 8 h with an exposure of 100 ms using a sCMOS camera (Orca Flash 4.0, Hamamatsu) with an effective pixel size of 65 nm.

### Machine learning

For each guide of the library, we computed a sgRNA activity score as the difference between its log2FC and the mean log2FC of all the guides within 6 Kbp, normalized between 0 and 1. The activity score of guides was computed as the mean activity from 4 independent assays measured in strain LC-E19 using the cosmid transduction assay.

After splitting the dataset into a training, validation and test sets, we implemented a neural network (Fig 3b) using Keras and TensorFlow. We used a model architecture inspired from our previous work (Cui et al. 2018) and consisting of 2 locally-connected layers of size 20 and 8, with kernel sizes of 4 and 7, followed by 2 dense layers of size 14 and 8. The layers sizes and kernel sizes were optimized to minimize the loss on the validation set using a grid search approach. We used the rectified linear unit (ReLU) activation function for all the neurons of these layers. Finally a densely connected single neuron predicted the sgRNA efficacy using a linear combination of the last dense layer. The network was trained to minimize the mean square error of the log2FC prediction with L2 regularization using the Adam optimizer (Kingma and Ba 2014). Training was interrupted when loss on the validation set ceased do decrease for more than two epochs.

To identify the positions used by the model to make its predictions we generated a set of 1000 random sequences, mutated each position *in silico*, and computed the effect of each mutation on the model prediction (Supplementary Figure 4). To measure the level of interaction between positions we generated all possible pairs of mutations for each sequence in a set of 200 random sequences, and compared the effect of individual mutations to that of pairs of mutations. Positions are interacting if the effect of a double mutation (Eij) is different from the sum of the effect of the single mutations (Ei+Ej). The heat map shows the average Euclidean distance between Eij and Ei+Ej for all pairs of positions (Figure 3).

### Model comparisons

The Fusi/Doench algorithm (from Doench 2016) was recoded from explanations provided in the article. In order to have a fair comparison with our model, all the features detailed in Doench 2016 were used except the cutting position and the amino acid percentage position. We then performed a 5-fold cross validation using a boosted regression tree on the FC+RES dataset from the same paper and obtained a Spearman coefficient similar to what was reported. In order to test the prediction capabilities of the Fusi/Doench algorithm on other datasets, we re-trained the algorithm on the whole FC+RES dataset and computed the correlation between the model predictions and the actual sgRNA activity.

Similarly, we recoded the Moreno-Mateos algorithm. Features far from the guide sequence that were used in the Moreno-Mateos were not kept as their value was not easily accessible for the FC+RES data. Removing them did not hurt the predictions. We then used a lassoCV regression and performed a 5-fold cross validation on the Moreno-Mateos dataset (obtained from Haeussler and colleagues (Haeussler et al. 2016)). We obtained similar prediction performance than described in their original paper (Spearman coefficient = 0.46). In order to test the prediction capabilities of the Moreno-Mateos algorithm on other datasets, we retrained the model on the whole Moreno-Mateos dataset and computed the correlation between the model predictions and the actual sgRNA activity.

## Data Access

The screen results are provided as Supplementary Dataset 1 and Supplementary Dataset 2. Other relevant data supporting the findings of the study are available in this article and its Supplementary Information files, or from the corresponding author upon request. An online version of the model is provided here: http://hub13.hosting.pasteur.fr:8080/CRISPRBact/

## Acknowledgements

We thank Gizem Ozbaykal for the help with the fluorescence and phase-contrast microscopy. DNA sequencing after Cas9 cleavage was performed by Cédric Fund from the Institut Pasteur Genomics Platform, a member of “France Génomique” consortium (ANR10-INBS-09-08). This work was supported by the European Research Council (ERC) under the Europe Union’s Horizon 2020 research and innovation program (grant agreement No [677823]); the French Government’s Investissement d’Avenir program; Laboratoire d’Excellence ‘Integrative Biology of Emerging Infectious Diseases’ [ANR-10-LABX-62-IBEID]; the Pasteur-Weizmann consortium, and a Ramon Areces Fellowship to B.G.

## Author contributions

B.G, L.C, J.W.N and D.B. designed the study and wrote the manuscript. B.G. and L.C performed the experiments. C.B. developed CRISPRbact. D.B. and J.W.N analysed the data. B.G, D.B. and J.W.N. wrote the manuscript.

